# Olfactory dysfunction in a novel model of prodromal Parkinson’s disease in adult zebrafish

**DOI:** 10.1101/2025.03.29.645796

**Authors:** Nathaniel W. Vorhees, Samantha L. Groenwold, Mackenzie T. Williams, Lexus S. Putt, Nereyda Sanchez-Gama, Grace A. Stalions, Gabriella M. Taylor, Heather E. Van Dort, Erika Calvo-Ochoa

## Abstract

Olfactory dysfunction is an early clinical marker of prodromal Parkinson’s disease (PD), yet the underlying mechanisms remain unclear. To explore this relationship, we developed a zebrafish model that recapitulates prodromal PD-associated olfactory impairment without affecting motor function. We used zebrafish, due to their olfactory system’s similarity to mammals and their unique nervous system regenerative capacity. By injecting 6-hydroxydopamine (6-OHDA) into the dorsal telencephalic ventricle, we observed a significant loss of dopaminergic (DA) periglomerular neurons in the olfactory bulb (OB) and retrograde degeneration of olfactory sensory neurons (OSNs) in the olfactory epithelium (OE). These alterations led to impaired responses to cadaverine, an aversive odorant, while responses to alanine, an attractive odorant, remained intact. 6-OHDA triggered robust neuroinflammatory responses that was attenuated by pranlukast, an anti-inflammatory drug. By 7 days post-injection, dopaminergic synapses in the OB were remodeled, OSNs in the OE appeared recovered, and neuroinflammation subsided, leading to full recovery of olfactory responses to cadaverine. These findings highlight zebrafish remarkable neuroplasticity and suggest this novel model of prodromal PD could provide valuable insights into early PD pathology. Understanding the interplay between dopaminergic loss, neuroinflammation, and olfactory dysfunction may inform therapeutic strategies for PD patients suffering from olfactory dysfunction.

## INTRODUCTION

Parkinson’s disease (PD) is a prevalent neurodegenerative disorder characterized by severe motor impairment, cognitive decline, and increased mortality in aging populations [1, 2]. Notably, over 95% of PD patients experience olfactory dysfunction, often preceding motor symptoms by several years, making it a key early clinical marker of prodromal PD [3-5].

The progressive and irreversible degeneration of dopaminergic (DA) neurons, particularly in the substantia nigra pars compacta, is a hallmark of PD, leading to widespread disruptions in dopaminergic neural circuits [6, 7]. Among the affected regions is the olfactory bulb (OB), the primary center for olfactory processing, which contains a dense population of dopaminergic periglomerular neurons crucial for odor detection and discrimination [8, 9]. Since the OB is one of the earliest structures impacted in PD [10, 11], dopaminergic dysfunction within this region may contribute to the olfactory deficits that are apparent years before motor symptoms manifest.

Although olfactory impairment in PD is extensive, research on the topic remains limited, highlighting the need for further investigation. To explore this, we used zebrafish, a well-established model for PD research due to its genetic similarity to humans (∼70% of genes are conserved; [12] and its remarkable capacity for nervous system regeneration [13].

Broadly, there are two types of experimental strategies to achieve PD-like pathology in zebrafish. Transgenic lines are used to study the genetic underpinnings of familial PD [14, 15], while neurotoxin-exposure models are used to replicate key features of sporadic PD, which accounts for ∼95% of cases (Tang, 2017). Among neurotoxins, the most commonly used include 6-OHDA, MPTP, rotenone, and paraquat, all of which induce mitochondrial dysfunction, oxidative stress, and neuroinflammation, ultimately leading to catecholaminergic neuronal loss [16-19]

We selected 6-OHDA for this study due to its specificity in targeting catecholaminergic neurons via uptake by the dopamine transporter, leading to localized neurodegeneration upon uptake [20]. Importantly, 6-OHDA injections allow for targeting of specific DA populations [21, 22] , making it an advantageous tool for targeting DA periglomerular neurons in the OB. Despite extensive research on PD in zebrafish, the relationship between dopaminergic loss and olfactory dysfunction in relationship with PD neuropathology has not been explored.

An important consideration is that olfactory assessments in zebrafish typically rely on motor-behavioral assays (kalueff), resulting in potential confounds if there are motor impairments, which is often the case with PD models. To overcome this challenge, we sought to develop a model of early PD-related olfactory dysfunction without locomotor deficits, recapitulating the prodromal stage of the disease. We achieved this by injecting 6-OHDA into the dorsal telencephalic ventricle of adult zebrafish, selectively targeting DA periglomerular neurons in the OB.

Here, we demonstrate that olfactory dysfunction results as a consequence of DA loss in the OB. Olfactory morphology and function recovered within seven days, underscoring the high neuroplasticity of the zebrafish olfactory system. This model of prodromal PD provides a valuable tool for investigating the early stages and progression of this neurodegenerative disease, and could offer new insights into the relationship between dopaminergic loss and olfactory dysfunction.

## RESULTS

### 6-OHDA injections in the dorsal telencephalic ventricle target dopaminergic neurons in the OB

We aimed to develop a model of olfactory dysfunction that mimics the prodromal stages of Parkinson’s disease (PD) without motor impairment. To achieve this, we injected 6-hydroxydopamine (6-OHDA) into the dorsal telencephalic ventricle of adult zebrafish, aiming to target dopaminergic (DA) periglomerular neurons in the olfactory bulb (OB) exclusively.

To assess the impact of 6-OHDA injections in the OB, we examined sagittal brain sections immunostained for tyrosine hydroxylase (TH), a marker of dopaminergic interneurons [23, 24]. At 1-day post-injection (dpi), we observed a significant loss of TH+ neuronal somata compared to controls (Figures 1A, B). However, no further decrease was detected at 3 or 7 dpi (Figures 1A, B), suggesting that 6-OHDA-induced DA neuron loss peaks at 1 dpi, consistent with previous reports [22].

**Figure 1.**
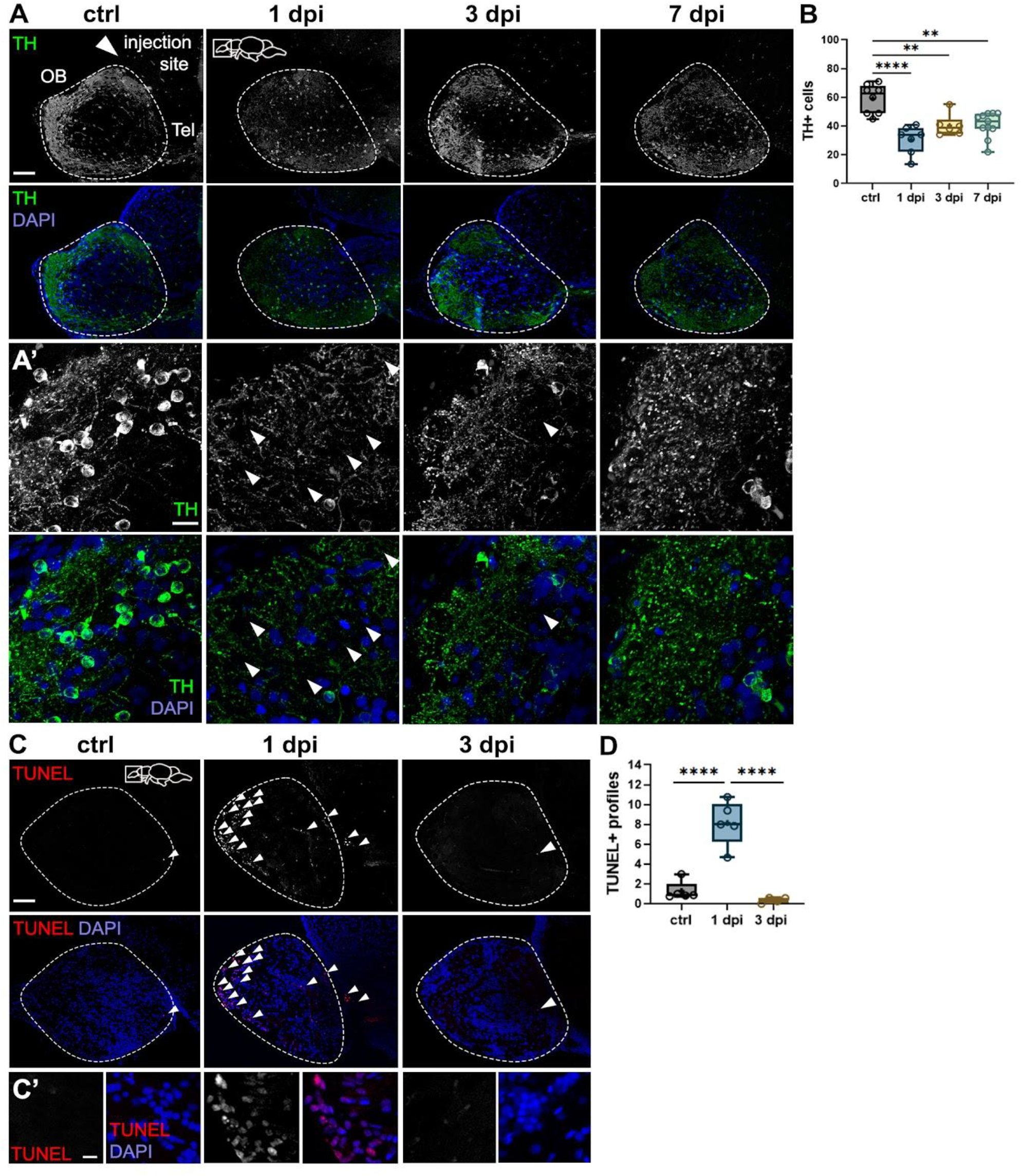
Effects of 6-OHDA injections on dopaminergic periglomerular neurons in the OB. (A) and (A’) Tyrosine hydroxylase (TH) immunohistochemistry of sagittal sections of the OB from control, 1-day post-injection (dpi), 3-dpi, and 7-dpi fish. 6-OHDA injection site at the dorsal telencephalic ventricle near the telencephalon (Tel) is shown with a white arrowhead in the top left panel. (A’) are magnified views of (A). Increased spacing in TH+ staining is indicated with white arrowheads. Green: TH; blue: DAPI. Scale bars: 100 μm in (A); 20 μm in (A’). (B) Quantification of the number of TH+ cells in sagittal sections of the OB from (A) (n = 6-11). ANOVA: F (4, 33) = 12.15, p < 0.0001. ctrl vs 1 dpi p < 0.0001, ctrl vs. 3 dpi p = 0.0050, ctrl vs. 7 dpi p = 0.0013. (C) (TdT) dUTP Nick-End Labeling (TUNEL) staining in sagittal sections of the OB from control, 1-dpi, and 3-dpi groups. TUNEL+ profiles are indicated with white arrowheads. Red: TUNEL; blue: DAPI. Scale bars: 100 μm in (C); 20 μm in (C’). (D) Quantification of average TUNEL+ profiles in OB sections from (C) (n = 4-5). ANOVA: F (2, 11) = 38.48, p < 0.0001. ctrl vs. 1 dpi p < 0.0001, 1 dpi vs. 3 dpi p < 0.0001. Box plots indicate mean (+), quartiles (boxes) and range (whiskers). One-way ANOVA, *p < 0.005, **p < 0.001, ***p = 0.0009, ****p < 0.0001.

High-magnification images revealed a notable disruption of DA neuron dendritic arbors in all injected groups compared to controls (Figure 1A’). In the 1-dpi group, TH+ puncta were more widely spaced across the parenchyma (arrowheads), indicating dendritic disorganization. However, by 7 dpi, the dendritic structure appeared closer to control levels than to 1 dpi (Figure 1A’), suggesting that the remaining periglomerular neurons in the OB exhibit dendritic plasticity in response to injury.

To determine whether 6-OHDA induces apoptosis, we performed a terminal deoxynucleotidyl transferase (TdT) dUTP Nick-End Labeling (TUNEL) assay. At 1 dpi, we observed a significant increase in TUNEL+ profiles in the OB glomerular layer, where periglomerular neurons reside (Figures 1C, D). By 3 dpi, TUNEL+ cell numbers returned to control levels (Figures 1C, D), indicating that apoptosis occurs rapidly and is not sustained [25]. This aligns with our finding that TH+ cell loss does not persist beyond 1 dpi (Figure 1B). These results demonstrate that 6-OHDA injections in the dorsal telencephalic ventricle effectively target DA periglomerular neurons in the OB, at least in part through apoptotic mechanisms.

### Periglomerular cell loss is associated with severe synaptic disruption in the OB

Olfactory sensory neurons (OSNs) form the first synapses of the olfactory pathway within spherical structures in OB parenchyma called glomeruli. These synapses, essential for odor processing, consist of OSN presynaptic termini and the dendrites of glutamatergic neurons (i.e., mitral and tufted cells; [26], and dopaminergic (DA) interneurons (i.e., periglomerular cells;[9]. Since 6-OHDA targets periglomerular cells and disrupts their dendritic arbors (Figure 1), we examined its effects on olfactory synaptic contacts. To do this, we qualitatively analyzed immunostained sections against the presynaptic marker synaptic vesicle protein 2 (SV2), which labels olfactory axon terminals, along with TH to visualize DA neurons. We focused on three large glomerular clusters apparent in sagittal OB sections: dorsal (dG), dorsolateral (dlG), and ventromedial (vmG) clusters (Figures 2A, B).

**Figure 2.**
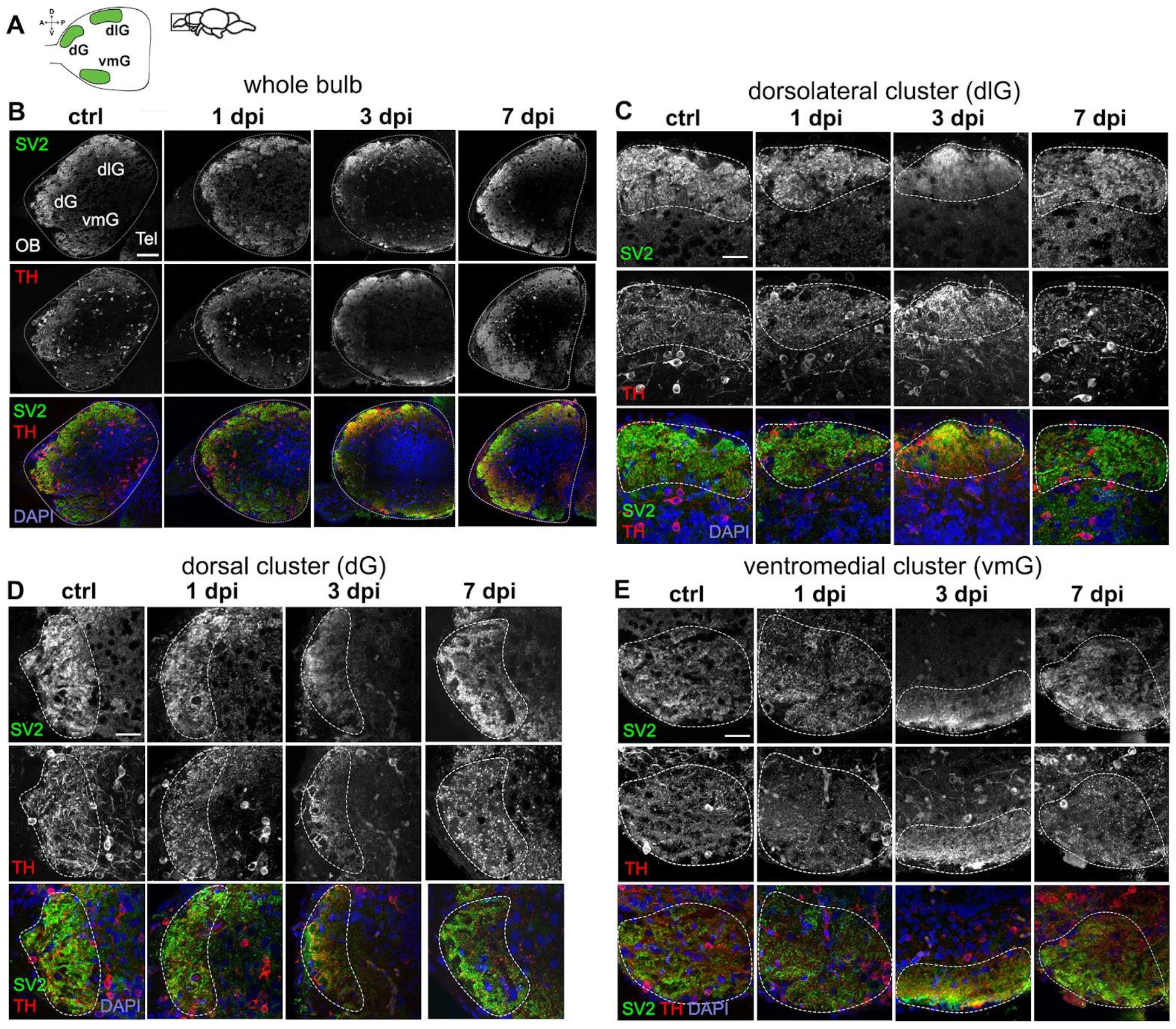
Dysregulation of synaptic contacts in the OB caused by 6-OHDA. (A) Schematic diagram of a sagittal view of the OB, indicating three glomerular clusters: dorsal (dG), dorsolateral (dlG), and ventromedial (vmG). (B-E) Double immunohistochemistry of synaptic vesicle protein 2 (SV2) and tyrosine hydroxylase (TH) in sagittal sections of the OB from controls, 1-dpi, 3-dpi, and 7-dpi groups. Whole bulb (B), and magnified views of: (C) dlG cluster, (D) dG cluster, and (E) vmG cluster. Green: Tbr2a; red: TH; blue: DAPI. Scale bars: 100 μm in (B); 50 μm in (C, D, E).

Individual glomeruli were identified based on SV2 staining (dotted lines in Figures 2C-E). We observed a pattern of synaptic disorganization followed by recovery across all glomeruli, indicating the widespread impact of 6-OHDA injections throughout the OB. In control fish, glomerular clusters appeared as distinct, round SV2-stained structures containing individual, smaller glomeruli [27]. TH-stained dendrites were present within these spaces – where they form synaptic contacts – while DA somata were located outside the cluster (Figures 2B, C).

At 1 dpi, TH+ puncta and somata showed clear disorganization, yet individual SV2-stained glomeruli remained distinguishable, albeit smaller than in controls (Figures 2B, C). By 3 dpi, glomerular organization was nearly lost, with no distinct glomeruli visible (Figures 2B, C). Both SV2 and TH staining were relocalized closer to the outer layers of the OB, suggesting synaptic decoupling and degeneration. By 7 dpi, most glomerular structures had recovered, resembling those of control fish (Figures 2B, C).

These findings suggest that while DA neuron loss peaks at 1 dpi, olfactory axon disorganization and synaptic degeneration occur more gradually over the following days. By 7 dpi, both periglomerular cell dendrites and olfactory presynaptic terminals appear remodeled and reorganized, highlighting the high plasticity of neurons within the olfactory system.

### Periglomerular and synaptic degeneration in the OB lead to impaired olfactory responses to selective odorants without altering locomotion

Our primary goal was to develop a model of PD-related olfactory dysfunction without motor impairment. To assess whether DA loss due to 6-OHDA injections leads to olfactory deficits, we employed olfactory-mediated behavioral assays. Additionally, we analyzed swimming parameters to determine if sham injections affect swimming performance. For this, we used two behavioral arenas to record responses to different odorants (Figures 3A, F). In the first experiment, we used a narrow rectangular tank to evaluate aversive responses to cadaverine, including freezing, darting, and sinking (Figure 3A).

**Figure 3.**
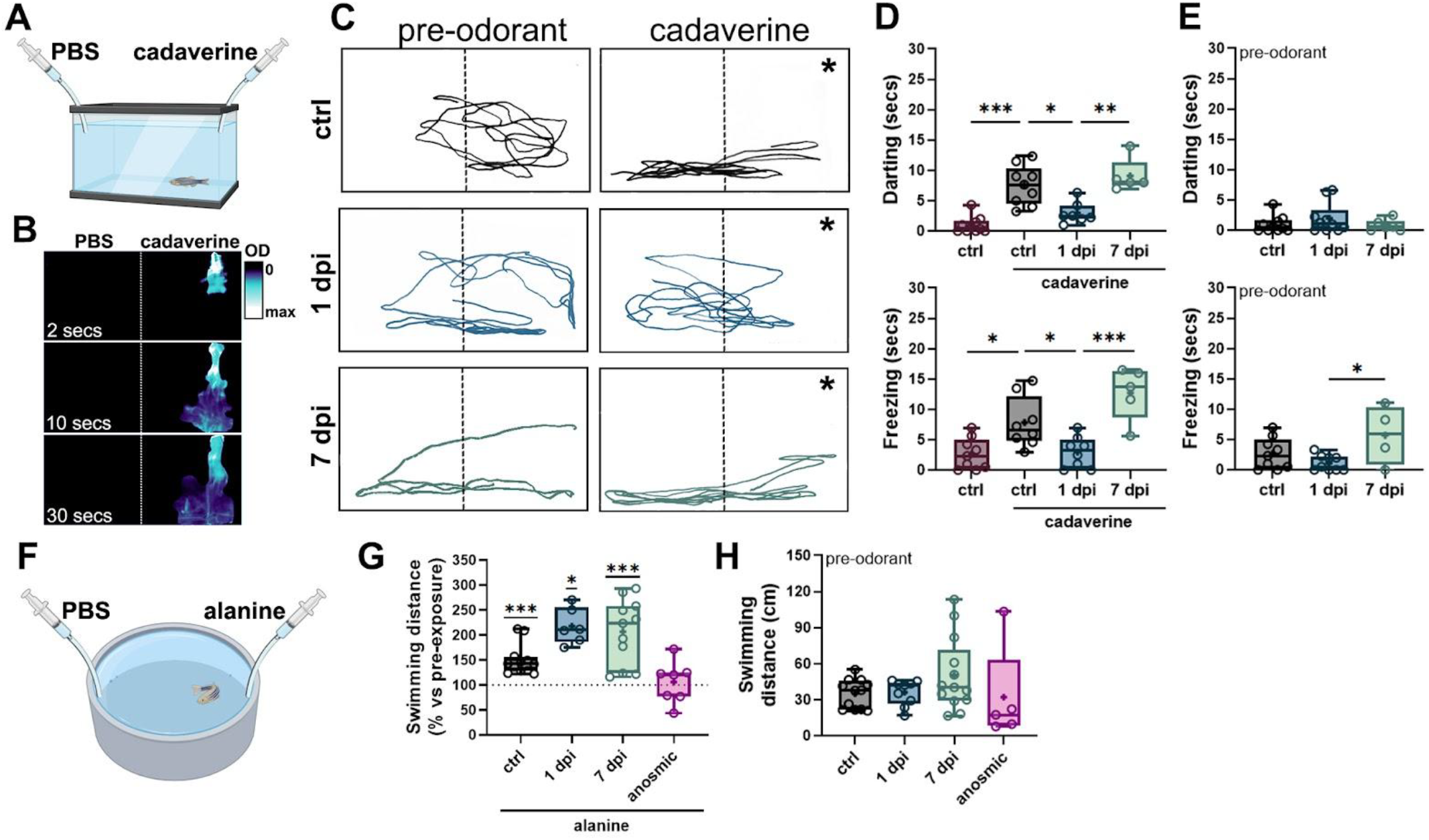
6-OHDA injections selectively reduce olfactory-mediated responses. (A) Schematic representation of the behavioral chamber used for studying responses to cadaverine. A rectangular, narrow chamber with a camera located in front was used to study fish’ vertical displacement following cadaverine exposure. (B) Time course of a dye distribution in the behavioral chamber from (A) indicating that odorant solutions remain in half of the chamber (indicated by a dotted line) during 30 seconds after cadaverine delivery. In (C), the asterisk (*) indicates the position where cadaverine solution was administered. (C) Representative swimming trajectories of zebrafish pre-(30 secs) and post-(30 secs) cadaverine delivery in controls (upper panels), 1-dpi (middle panels), and 7-dpi (lower panels) fish. (D) Quantification of swimming responses to cadaverine in controls, 1-dpi, 3-dpi, and 7-dpi (n=5-10). Top panel: time spent darting after cadaverine exposure. ANOVA F (3, 27) = 19.93, p < 0.0001. ctrl vs. control cad p < 0.0001, ctrl cad vs. 1 dpi p = 0.0033, 1 dpi vs. 7 dpi p = 0.0007. Bottom panel: time spent freezing after cadaverine exposure. ANOVA F (3, 26) = 12.28, p < 0.0001. c trl vs. control + cad p = 0.0206, ctrl cad vs. 1 dpi p = 0.0379, 1 dpi vs. 7 dpi p = 0.0002. (E) Quantification of darting (top panel) and freezing (bottom panel) before odorant exposure (n=4-9). For freezing: ANOVA F (2, 19) = 4.088, p = 0.0334. 1 dpi vs. 7 dpi p = 0.0261. (F) Schematic representation of the behavioral chamber used for studying responses to alanine. A larger circular chamber with an overhead camera was used to study fish’ swimming distance following alanine exposure. (G) Quantification of swimming responses to alanine compared against pre-odorant exposure in controls, 1-dpi, 3-dpi, 7-dpi, and anosmic fish (n = 6-12). One-sample Wilcoxon test when compared to baseline of 100% (no change). ctrl = p = 0.0005, 1 dpi p = 0.0312, 7 dpi p = 0.0010. (H) Quantification of swimming responses before odorant exposure (n = 6-12). Box plots indicate mean (+), quartiles (boxes) and range (whiskers). *p < 0.005, **p < 0.001, ***p = 0.0009, ****p < 0.0001.

Individual fish were recorded for 30 seconds before (pre-odorant) and after cadaverine (cad) exposure. The odorant remained confined to one half of the chamber (Figure 3B), allowing us to assess aversive swimming responses (Figure 3C). Control fish exhibited typical olfactory-mediated avoidance, moving toward the non-cadaverine side of the tank and remaining in the bottom (Figure 3C, top panel, cadaverine side indicated with an asterisk). They also displayed significant increases in darting (Figure 3D, top panel) and freezing (Figure 3D, bottom panel) compared to non-exposed controls. In contrast, 1-dpi fish maintained similar swimming patterns after odorant exposure (Figure 3C, middle panel) and did not exhibit erratic swimming or freezing to cadaverine (Figure 3D), indicating an impaired olfactory response to this odorant.

Since extensive synaptic remodeling was observed in the OB by 7 dpi, we tested whether olfactory function could recover. In line with our predictions, 7-dpi fish exhibited swimming trajectories and behaviors similar to controls (Figure 3C, bottom panel), including increased darting and freezing (Figure 3D), significantly different from the 1-dpi group, suggesting that olfactory responses to cadaverine are restored by 7 dpi. To determine whether 6-OHDA altered baseline darting and freezing behaviors, we analyzed pre-odorant trial data. 1-dpi fish showed no differences in darting or freezing compared to controls (Figure 3E). However, in the 7-dpi group, freezing time was significantly increased compared to the 1-dpi group, but not to controls (Figure 3E).

To assess whether olfactory dysfunction was more widespread, we used an open swimming arena to assess olfactory responses to alanine, an amino acid that elicits attraction and appetitive behaviors, reflected as increased swimming distance and speed [28] (Figure 3F). As expected, we found that control fish responded to alanine by significantly increasing the swimming distance after alanine exposure, compared to a 100% baseline (vs. non-exposed fish, Figure 3G). Both 1-dpi and 7-dpi fish exhibited similar responses to alanine, suggesting that olfactory responses to this odorant remained intact (Figure 3G). Since alanine is palatable for some teleost species [29], we investigated whether the observed behavioral responses could be attributed to gustation. For this, we generated a group of anosmic fish by occluding the nares with surgical adhesive. These fish did not increase their swimming distance following alanine exposure (Figure 3G), confirming that the responses to alanine observed were olfactory-mediated.

To assess potential swimming impairment due to 6-OHDA injection, we analyzed pre-odorant swimming distances across all groups. No significant differences were found. (Figure 3H). These findings indicate that 6-OHDA injections do not impair olfactory responses to alanine or affect swimming performance in an open arena. Overall, our results demonstrate that 6-OHDA injections selectively disrupt olfactory responses. While fish lose the ability to detect cadaverine, their response to alanine remains intact. Notably, cadaverine sensitivity is restored by 7 dpi. Additionally, 6-OHDA injections do not impair swimming ability at the tested timepoints. Together, these findings confirm that our model successfully replicates prodromal PD with olfactory dysfunction in the absence of motor impairment.

### Astroglial cells are activated in the OB following 6-OHDA injections

Neuroinflammation is a hallmark of Parkinson’s disease (PD) pathology, with sustained astroglial and microglial activation contributing to synaptic and neuronal degeneration [30, 31]. In contrast, zebrafish exhibit a rapid resolution of post-injury neuroinflammatory responses without forming a permanent astroglial scar [32], which has been shown to hinder repair in mammalian brains [33]. Thus, we sought to determine whether 6-OHDA ventricular injections induce astroglial activation in the OB.

To investigate this, we analyzed OB tissue immunostained against the astroglial marker, glial fibrillary acidic protein (GFAP). Consistent with previous reports, GFAP staining was observed in the olfactory nerve (ON) and the outermost OB layer, the olfactory nerve layer (ONL), both of which are composed of olfactory axons stemming from the OSNs from the olfactory epithelium (OE) (Figure 4A, arrowheads) [34]. In 6-OHDA-injected fish, we detected a significant increase in GFAP expression at all timepoints, peaking at 3 dpi (Figures 4A, B). To determine whether this astroglial activation resulted from 6-OHDA or the injection procedure itself, we co-injected 6-OHDA with the anti-inflammatory drug pranlukast (PRAN) and examined the tissue at 1 dpi. Our findings revealed no significant differences between the control and PRAN-treated groups (Figures 4A, B), indicating that the 6-OHDA-mediated inflammatory response can be attenuated with anti-inflammatory treatment.

**Figure 4.**
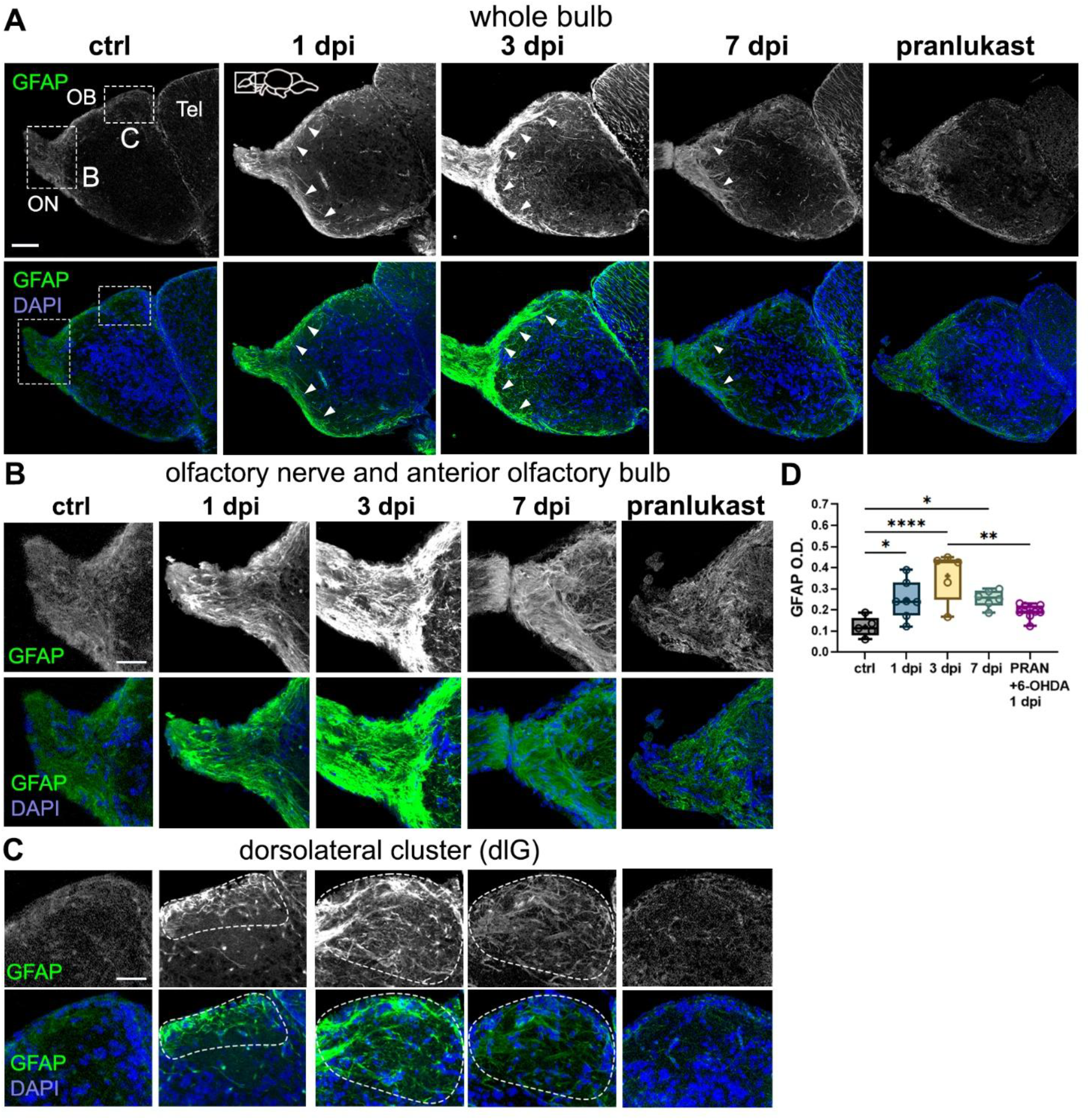
6-OHDA injections lead to astroglial activation in the OB. (A-C) Glial fibrillary acidic protein (GFAP) immunohistochemistry in sagittal sections of the OB from controls, 1-dpi, 3-dpi, 7-dpi, and 6-OHDA with the anti-inflammatory drug pranlukast (PRAN). Whole bulb (A) and magnified views of: (B) the olfactory nerve and anterior OB, and (C) the dorsolateral (dlG) glomerular cluster (indicated by dotted lines). Green: GFAP; blue: DAPI. Scale bars: 100 μm in (A); 50 μm in (B, C). (D) Quantification of GFAP O.D. in OB sections from (A) (n = 5-9). ANOVA: F (4, 27) = 8.478, p = 0.0001. ctrl vs. 1 dpi p = 0.0306, ctrl vs. 3 dpi p < 0.0001, ctrl vs. 7 dpi p = 0.0275. 1 dpi vs. 3 dpi p < 0.0001. 3 dpi vs. PRAN p = 0.0017. Box plots indicate mean (+), quartiles (boxes) and range (whiskers). One-way ANOVA, *p < 0.005, **p < 0.001, ***p = 0.0009, ****p < 0.0001.

Further analysis confirmed a marked increase in GFAP+ staining and branching across the ON and ONL in all injected groups, except for the PRAN-treated group (Figure 4B). We observed a dramatic increase of GFAP+ processes surrounding the dorsolateral glomerulus (dlG, dotted line), particularly at 3 dpi, where astroglial processes distinctly outlined the glomerular cluster. This pattern was absent in the control and PRAN groups (Figure 4C). Interestingly, the peak in astroglial activation at 3 dpi coincides with the timepoint at which the synaptic disorganization is the most pronounced (Figure 2), suggesting a potential role for astroglia in synaptic remodeling.

### 6-OHDA promotes microglial activation and leukocyte migration to the OB

A key component of the neuroinflammatory response, which is also heightened in PD, is microglial activation [35]. To further investigate the neuroimmune response to 6-OHDA injections, we performed histological characterization using the leukocyte/microglial marker lymphocyte cytosolic protein 1 (Lcp1, also known as L-plastin). Lcp1 is primarily expressed in microglia within the central nervous system (CNS) and in peripheral leukocytes. Under normal physiological conditions, peripheral immune cells are scarce in the CNS, making Lcp1 a *bona fide* microglial marker [36].

In control fish, we observed sparse Lcp1+ microglial cells distributed across the OB, with most cells displaying a stellate morphology characteristic of a resting state (Figures 5A, B) [37]. However, in 1-dpi and 3-dpi fish, there was a significant increase in Lcp1+ cells in the OB, suggesting an infiltration of peripheral leukocytes (Figure 5A, F). Higher magnification images of the OB periphery (Figure 5C) and of the interphase between the telencephalon and OB (Figure 5D) confirmed a noticeable increase in Lcp1+ cells at these timepoints, indicative of cell infiltration. Consistent with our earlier findings of astroglial activation around glomerular clusters, particularly in the dorsolateral glomerulus (dlG) (Figure 4C), we also observed an accumulation of Lcp1+ cells in this region at 1 dpi and 3 dpi (Figure 5E).

**Figure 5.**
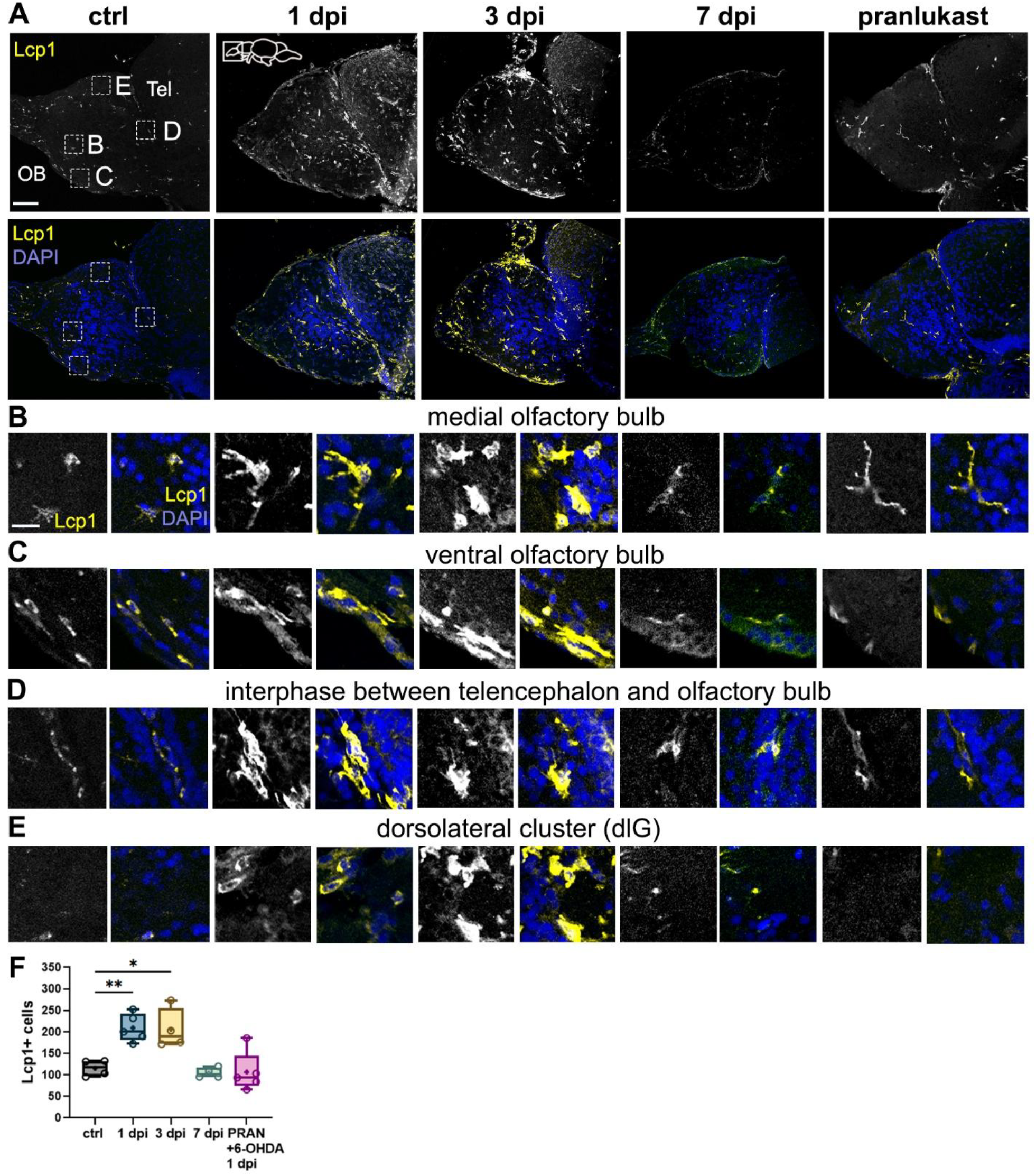
Microglial activation and leukocyte migration in the OB following 6-OHDA injection. (A-E) Lymphocyte cytosolic protein 1 (Lcp1) immunohistochemistry in sagittal sections of the OB from controls, 1-dpi, 3-dpi, 7-dpi, and 6-OHDA with the anti-inflammatory drug pranlukast (PRAN). Whole bulb (A) and magnified views of: (B) middle region of the OB, (C) dlG cluster region, (D) ventral region, and (E) the interphase between the telencephalon and the OB. Yellow: Lcp1; blue: DAPI. Scale bars: 100 μm in (A); 20 μm in (B-E). (F) Quantification of Lcp1+ cells in OB sections from (A) (n = 4-5). ANOVA: F (4, 17) = 10.66, p = 0.0002. ctrl vs. 1 dpi p = 0.0074, ctrl vs. 3 dpi p = 0.0151, ctrl vs. 7 dpi p = 0.0028, 1 dpi vs. 7 dpi p = 0.0028, 1 dpi vs. PRAN p = 0.0019, 3 dpi vs. 7 dpi p = 0.0061, 3 dpi vs. PRAN p = 0.0046. Box plots indicate mean (+), quartiles (boxes) and range (whiskers). One-way ANOVA, *p < 0.005, **p < 0.001, ***p = 0.0009, ****p < 0.0001.

In addition to increased cell migration, we noted striking changes in the size and morphology of Lcp1+ cells in the 1-dpi and 3-dpi groups, indicating activation states. These alterations, including larger somata with extensive processes (indicative of migration) and rounded, enlarged somata (suggestive of phagocytic activity), were most pronounced at 3 dpi (Figures 5B–E). Interestingly, by 7 dpi, the leukocytic/microglial response had largely subsided (Figures 5A, B), and this response was notably hampered in fish co-treated with PRAN (Figures 5A, B). Taken together, these findings demonstrate that 6-OHDA injections trigger robust microglial and leukocytic activation and migration in the OB, a response that resolves by 7 dpi. Notably, the peak activation and phagocytic activity of Lcp1+ cells at 3 dpi align with the observed increase in astrocytic activity at this same time point (Figure 4), suggesting that astroglial and microglial cells work collaboratively at axonal and synaptic sites to facilitate their remodeling and recovery.

### 6-OHDA-mediated changes in the OB cause retrograde degeneration and cell proliferation in the OE

Given the bidirectional communication between the peripheral olfactory epithelium (OE) and the olfactory bulb (OB), we aimed to determine whether the neuronal and synaptic alterations induced by 6-OHDA affected the OE through retrograde degeneration mechanisms [38, 39]. To assess this, we analyzed sections of the olfactory organ immunostained for HuC/D, a pan-neuronal marker in zebrafish. The olfactory organ comprises both sensory (HuC/D+) and non-sensory epithelium, (delineated in dotted lines, Figure 6A). We observed a significant reduction in HuC/D staining in the 1-dpi and 3-dpi groups compared to controls (Figures 6A, A’, B), suggesting olfactory sensory neuron (OSN) loss or dysfunction in the OE. However, there were no noticeable changes in the overall size or gross morphology of the olfactory organ (Figure 6A). By 7 dpi, HuC/D staining had returned to control levels, indicating OSN recovery (Figures 6A, A’, B).

**Figure 6.**
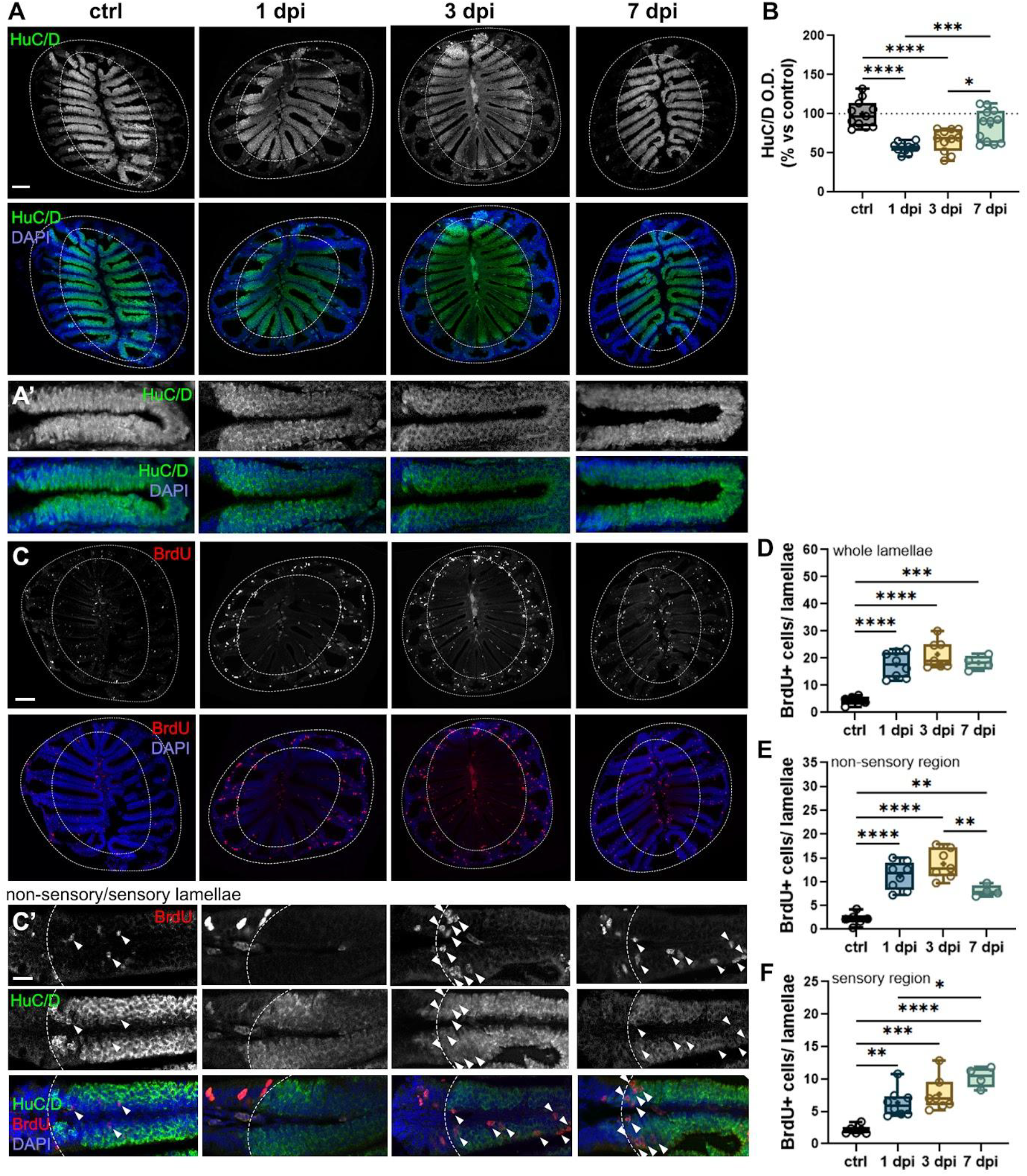
6-OHDA injections in the OB cause retrograde degeneration in the OE. (A, A’) HuC/D immunohistochemistry of OE sections from controls, 1-dpi, 3-dpi, and 7-dpi groups. (A’) are magnified views of the OSN-containing sensory lamellae (HuC/D+) from (A). Scale bars: 100 μm in (A); 20 μm in (A’). (B) Quantification of percent change (vs. control) in optical density (O.D.) of HuC/D in the OE from (A) (n = 11-12). ANOVA F (3, 41) = 16.78, p < 0.0001. ctrl vs. 1 dpi p < 0.0001, ctrl vs. 3 dpi p < 0.0001, 1 dpi vs. 7 dpi p = 0.0005, 3 dpi vs. 7 dpi p = 0.0286. (C, C’) Double immunohistochemistry of BrdU and HuC/D in sections of the OE of controls, 1-dpi, 3-dpi, and 7-dpi fish. (C’) Magnified views of (C), showing the sensory and non-sensory regions of the OE lamellae (indicated by dashed lines and HuC/D staining). BrdU+ cells found in the sensory area (HuC/D+) are indicated with white arrowheads. Red: BrdU; green: HuC/D; blue: DAPI. Scale bars: 100 μm in (C); 20 μm in (C’). (D)Quantification of BrdU+ cells in whole lamellae from (A) (n=4-8). ANOVA F (3, 21) = 21.26, p < 0.0001. ctrl vs. 1 dpi p < 0.0001, ctrl vs. 3 dpi p < 0.0001, ctrl vs. 7 dpi p = 0.0002. (E) Quantification of BrdU+ cells in the non-sensory lamellae from (A) (n=4-8). ANOVA F (3, 21) = 25.83, p < 0.0001. ctrl vs. 1 dpi p < 0.0001, ctrl vs. 3 dpi p < 0.0001, ctrl vs. 7 dpi p = 0.0073, 3 dpi vs. 7 dpi p = 0.0075. (F) Quantification of BrdU+ cells in the sensory lamellae (HuC/D+) from (A) (n=4-8). ANOVA F (3, 21) = 14.86, p < 0.0001. ctrl vs. 1 dpi p = 0.0046, ctrl vs. 3 dpi p = 0.0004, ctrl vs. 7 dpi p < 0.0001, 3 dpi vs. 7 dpi p = 0.0075, 1 dpi vs. 7 dpi p = 0.0205. Box plots indicate mean (+), quartiles (boxes) and range (whiskers). One-way ANOVA, *p < 0.005, **p < 0.001, ***p = 0.0009, ****p < 0.0001.

The OE is known for its rapid regenerative capacity following various types of damage[40]. Given that OSNs recovered by 7 dpi, we hypothesized that neurogenesis was upregulated following the injection, thus facilitating OE repair. To test this, we labeled newborn cells generated within the first 24 hours after 6-OHDA injection with bromodeoxyuridine (BrdU) and tracked them at different timepoints. Our results revealed a significant increase in BrdU+ profiles across the entire lamellae in all injected groups compared to controls (Figures 6C, D).

The OE contains two main progenitor populations: globose basal cells (GBCs), which reside in the lateral, non-sensory regions of the OE and are constitutively proliferative; and horizontal basal cells (HBCs), located along the basal lamina of the sensory epithelium (Figure 6C’, delineated by dotted lines). Under normal conditions, HBCs remain quiescent but become activated in response to epithelial damage [41]. By analyzing the localization of BrdU+ cells within the epithelium, we assessed neurogenesis, as neurons derived from progenitors migrate toward the sensory lamellae, typically positioned more apically than HBCs, which remain near the basal lamina.

Our findings showed an increase in BrdU+ cells in both the non-sensory and sensory regions in all injected groups compared to controls, suggesting activation of both types of progenitors (Figures 6C-F). Notably, the number of BrdU+ cells in the non-sensory epithelium exhibited a decreasing trend in the 7-dpi group, as indicated by a significant difference compared to the 1-dpi group (Figure 6E). Conversely, in the sensory epithelium, we observed the opposite trend: while all injected groups displayed a significant increase in BrdU+ cells compared to controls (Figures 6C’, arrowheads; F), BrdU+ profiles were further elevated at 7 dpi, as indicated by a significant increase relative to the 1-dpi group (Figure 6F). Together, these results indicate that 6-OHDA injections lead to retrograde degeneration of OSNs in the OE, followed by recovery by 7 dpi. This process is accompanied by increased cell proliferation and neurogenesis in the OE, particularly evident at 7 dpi — coinciding with the restoration of olfactory axonal synaptic morphology and olfactory function (Figures 2, 3).

### Peripheral leukocytes migrate to OE’s sensory epithelium following 6-OHDA injections

We hypothesized that the observed alterations in the OE would be accompanied by an increase in inflammatory processes. To investigate this, we performed Lcp1 staining in OE sections to assess the presence of peripheral leukocytes in the OE. In control fish, we identified a resident population of Lcp1+ cells primarily localized within the non-sensory lamellae, with only a few individual cells dispersed throughout the sensory epithelium (Figure 7A), consistent with previous reports describing a similar organization of neutrophils (the most abundant leukocyte) in the OE [42].

**Figure 7.**
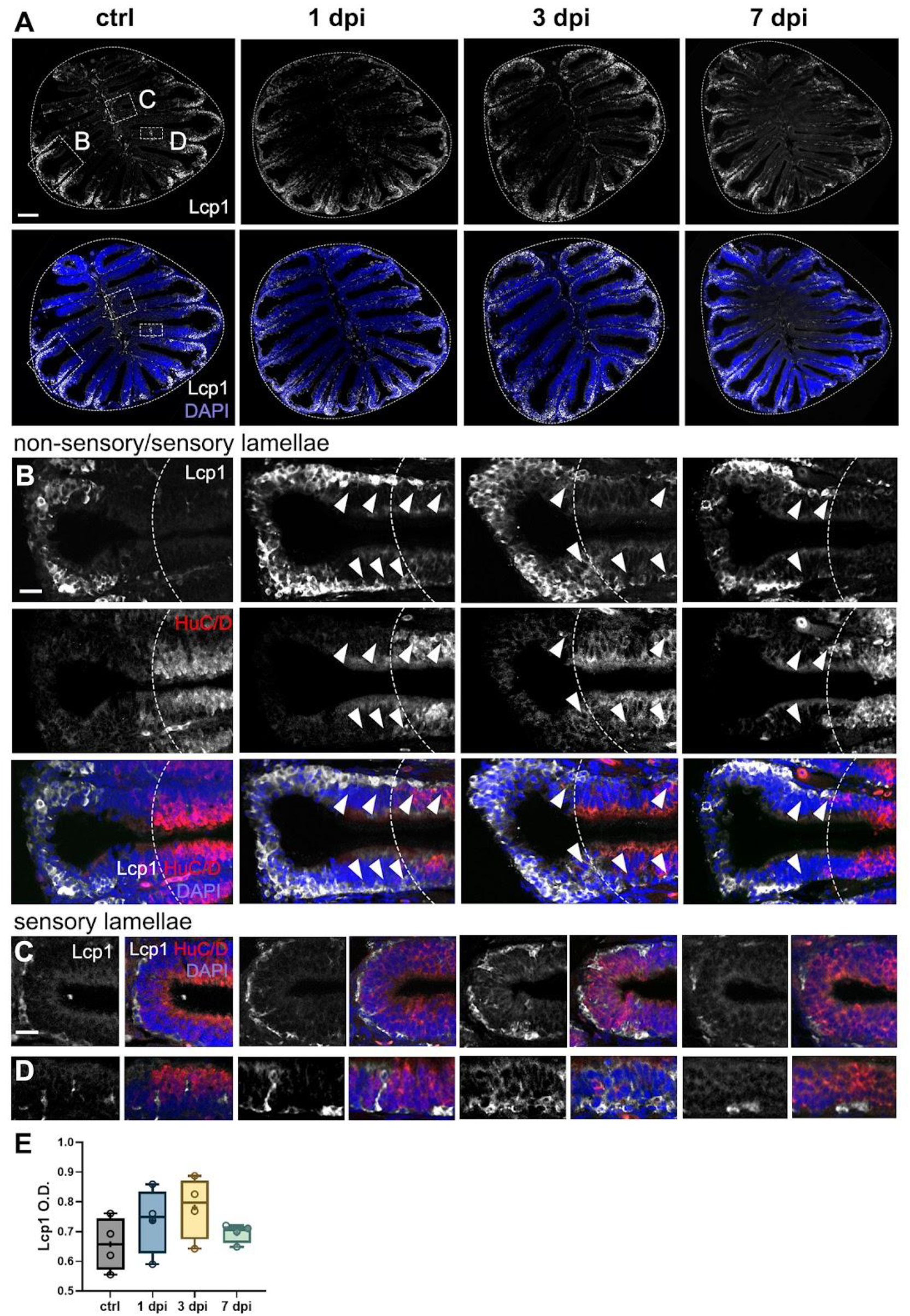
Leukocytic migration within the OE caused by 6-OHDA injections in the OB. (A-D) Lymphocyte cytosolic protein 1 (Lcp1) immunohistochemistry in sections of the OE from controls, 1-dpi, 3-dpi, and 7-dpi. Whole OE (A) and magnified views of: (B) distal epithelial fold region, where the OSN-containing sensory region is indicated by HuC/D staining and a dotted line; (C) medial sensory region (HuC/D+) adjacent to the central raphe; and (D) sensory epithelium (HuC/D+). White: Lcp1; blue: DAPI. Scale bars: 100 μm in (A); 20 μm in (B-D). (E) Quantification of Lcp1+ cells in OE sections from (A) (n = 4). ANOVA: F (3, 12) = 1.42, p = 0.2840. Box plots indicate mean (+), quartiles (boxes) and range (whiskers).

Interestingly, we did not detect significant differences in Lcp1+ optical density (O.D.) across groups (Figures 7A, E). We assessed O.D. as the high density of Lcp1+ cells in the OE precluded reliable individual cell counts. Upon closer examination of magnified views of the epithelia at the interphase between the non-sensory and sensory regions (Figure 7B, dotted lines) revealed that in the 1-dpi and 3-dpi groups, Lcp1+ cells migrated toward the HuC/D+ sensory region

(Figure 7B, arrowheads). By 7 dpi, the distribution of leukocytes resembled that of control fish, with most cells residing in the non-sensory epithelium (Figure 7B). We also examined the medial epithelial folds adjacent to the central raphe, a known site of immune cell infiltration into the OE [42, 43]. At 1 dpi and 3 dpi, we observed an increased presence of Lcp1+ cells lining the basal end of the epithelium near the raphe, suggestive of cell migration, but this increase was not evident at 7 dpi (Figure 7C). Additionally, in the 1-dpi and 3-dpi groups, leukocytes were embedded throughout the width of the sensory epithelium, suggesting that these immune cells migrate both horizontally and apically within the sensory lamellae (Figure 7D). Collectively, these findings indicate that 6-OHDA injections in the OE lead to the redistribution of resident Lcp1+ leukocytes, shifting from the non-sensory lamellae to the OSN-rich sensory regions.

## DISCUSSION

In this study, we developed a novel model of prodromal Parkinson’s disease (PD) characterized by olfactory impairment without motor deficits. To this end, we injected 6-OHDA into the dorsal telencephalic ventricle to target dopaminergic (DA) periglomerular neurons in the olfactory bulb (OB) in adult zebrafish. This approach enabled us to investigate the early effects of dopaminergic loss in the olfactory system in a model that recapitulates some pathological features of early PD.

### 6-OHDA-mediated dopaminergic loss in the OB leads to olfactory dysfunction

Our findings reveal that 6-OHDA injections led to a decreased olfactory-mediated response to cadaverine, an odorant that elicits avoidance behaviors, while responses to alanine, an odorant prompting attraction and appetitive behaviors, remained unchanged. This selective olfactory impairment suggests that certain olfactory circuits may be more vulnerable to dopaminergic loss than others, possibly reflecting differential susceptibility among olfactory sensory neuron (OSN) subtypes. Given that amino acids serve as cues for detecting food sources, the teleost olfactory system has adapted to prioritize amino acid detection, with a greater proportion of OSNs dedicated to processing these stimuli compared to aversive odorants [44, 45]. Cadaverine is recognized by a subset of ciliated OSNs that extend projections to a single glomerulus in the dorsolateral cluster (dlG, [45, 46], while amino acids, such as alanine, activate a larger group of microvillous OSN that project to lateral glomeruli [47, 48].

Importantly, our results confirm prior findings by Godoy et al., who reported that DA neuron ablation in the OB leads to impaired responses to cadaverine [49]. Although there are important differences in the onset of olfactory dysfunction, in both studies this dysfunction correlates with the peak loss of periglomerular neurons. Godoy et al., employed an elegant chemogenic strategy to ablate DA neurons selectively, which led to peak DA neuron loss and olfactory dysfunction at 7 days post-ablation. We observed olfactory dysfunction as early as 1 day after 6-OHDA exposure, coinciding with the peak DA neuron loss in our model. This discrepancy may be due to the neurotoxic effects of 6-OHDA, which ablates DA neurons more rapidly.

Dopaminergic neuron regeneration in the OB is well documented in zebrafish and other species [22, 49, 50], and is thought to contribute to the recovery of olfactory function following DA loss [21, 49]. Our data show that DA neuron numbers at 7 dpi remain unchanged from 1 dpi, indicating that no new neurons have yet been incorporated into the OB. This aligns with previous reports indicating that bulbar neuron regeneration occurs at least two weeks after neuronal birth [22, 39, 49, 51]

Thus, an important contribution of this work is that olfactory recovery occurred independently of periglomerular neuron regeneration, instead coinciding with compensatory synaptic remodeling in the OB, suggesting that plasticity mechanisms contribute to functional restoration before newly generated DA neurons integrate into OB circuits. Future studies should assess whether DA neuron numbers eventually recover in this model of neurotoxicity and, if so, the precise timeline of neuronal replacement.

### Neuroinflammatory processes parallel synaptic disruption and recovery in the OB

Our results reveal a time-dependent disruption and recovery of synaptic architecture in the OB that mirrors neuroinflammatory responses. Although periglomerular neuron loss peaks at 1 dpi, synaptic dysregulation is most pronounced at 3 dpi, coinciding with increased astroglial and microglial activation. This aligns with previous findings that demonstrate that neuroinflammatory responses in the zebrafish brain play a critical role in repair following neural injury [52, 53].

Unlike mammals, zebrafish lack stereotypical stellate GFAP+ astrocytes [54]. In lieu of these astrocytes, GFAP+ cells have similar morphologies and properties as radial glia in regions like the telencephalon and the optic tectum [55, 56]. In the olfactory system, GFAP+ astroglial cells have been described as olfactory ensheathing cells (OECs), which envelop olfactory axons from the OE basal lamina to the OB glomerular layer, where axonal termini are located [34]. OECs are unique glial cells that share properties with astrocytes and oligodendrocytes [57, 58] as they secrete neurotrophic factors that support axonal survival, growth, and remyelination [59].

A noteworthy feature of the glial response was the enlargement and dynamic nature of GFAP+ processes, which clearly delineated glomerular sites throughout the post-injection period. This suggests that OECs may contribute to synaptic remodeling, possibly through phagocytosis of damaged synapses and/or secretion of neurotrophic factors [58]. Another important role of OECs is to guide immature axons to their OB targets [60]. We demonstrate that 6-OHDA injections stimulate neurogenesis in the OE, likely as a consequence of decreased OSN density. It is possible that activated OECs facilitate proper axonal targeting of newborn OSNs, contributing to the synaptic remodeling observed at 7 dpi.

Furthermore, we observed pronounced microglial activation and leukocyte recruitment in the OB at 1 and 3 dpi, coinciding with peak astroglial responses, supporting the idea of a coordinated neuroimmune response. Under physiological conditions, microglia are the sole resident immune cells in the central nervous system (CNS), where they continuously survey the environment and facilitate homeostasis. In control fish, microglia were sparse and presented a ramified morphology. Following 6-OHDA injections, we report increased numbers of Lcp1+ cells in the OB, indicating that peripheral leukocytes infiltrated the CNS [61]. Accompanying the increase in the number of Lcp1+ cells in the OB, we observed noticeable changes in cell morphology, indicating infiltration and activation [37]. Leukocyte migration was the most evident in the periphery of the OB as well as in the interphase between the telencephalon and the OB, supporting previous reports of these cells infiltrating the brain through blood vessels and ventricles [61]. Infiltrating leukocytes at these locations exhibited a transitioning, motile state, characterized by an enlarged soma with shorter processes.

Activated microglia adopted an ameboid morphology, characterized by large, rounded somata with retracted processes, reflecting a phagocytic state. This morphology closely resembles that of infiltrating macrophages, which aid microglia in debris clearance and secretion of cytokines that modulate the neuroinflammatory response [37, 62]. At 3 dpi, we observed an increase of phagocytic Lcp1+ cells at 3 dpi, particularly around the olfactory nerve layer and glomerular clusters, suggesting that active phagocytosis of extracellular and axonal debris contributes to synaptic remodeling. Coupled with our observations of concomitant astroglial responses, we propose that OECs and microglia act synergistically to facilitate recovery and the establishment of new synaptic contacts following periglomerular cell loss. Notably, co-administration of the anti-inflammatory drug pranlukast mitigated both astrocytic and microglial activation, indicating that the inflammatory response mediated by 6-OHDA can be hampered by anti-inflammatory treatment, indicating that OB inflammation is due to 6-OHDA’s neurotoxic insult.

### 6-OHDA i.c.v. injections lead to retrograde degeneration and neurogenesis in the peripheral OE

Another major finding of this study is the evidence of retrograde degeneration of OSNs in the OE, indicated by a significant reduction of HuC/D+ OSNs at 1 dpi and 3 dpi, followed by recovery of neuronal density. These results are consistent with prior reports that neurodegeneration in the OB or in the ON lead to retrograde degeneration [38, 39, 63], underscoring the bidirectional communication between the peripheral and central components of the olfactory system. Our findings suggest that early olfactory deficits in our model likely result from both central and peripheral neuronal degeneration.

OSN dysfunction triggered local cell proliferation and neurogenesis, as indicated by BrdU labeling. Notably, the increase of BrdU+ profiles in the sensory region of the OE lamellae, coincides with the recovery of cadaverine-induced behavioral responses at 7 dpi. Since DA neurons in the OB had not recovered by this timepoint, we propose that recovery of OSN integrity coupled with synaptic remodeling in glomerular sites contribute to olfactory function recovery. Furthermore, the redistribution of leukocytes to the sensory region of the OE following 6-OHDA injections suggest that peripheral immune cells contribute to OSN repair and regeneration, echoing the response of their counterparts in the OB.

In conclusion, we describe a novel model of olfactory dysfunction associated with early PD pathology in adult zebrafish. Our results suggest that early-stage olfactory dysfunction in PD may arise from a combination of dopaminergic loss and synaptic dysregulation in the OB and retrograde degeneration of OSNs in the OE. We report that olfactory dysfunction and synaptic degeneration were recovered by 7 dpi, highlighting the remarkable and unique regenerative abilities of zebrafish. Future studies should explore the molecular mechanisms underlying this recovery, including the role of neurotrophic signaling and immune-mediated repair. Our findings underscore the use of zebrafish as an amenable model for investigating early PD-related olfactory dysfunction, with the potential for the development of recovery interventions for PD patients suffering from olfactory loss.

## ACKNOWLEDGEMENTS

We are grateful to current and past members of the Calvo lab for support throughout the completion of these studies; in particular to Theodore Lockett, Helen Dodge, Sarah Heinowski, Nathan Koorndyk, and Mackenzie Korff for technical support. We acknowledge the following institutions and agencies that have generously funded this work. ECO was supported by: the Kenneth Campbell Foundation, the International Brain Research Organization (IBRO, Rising Stars Award), and Hope College.

## Conflict of interest

The authors declare no competing financial interests.

## Author contributions

NWV, SLG, and ECO developed the i.c.v. model. NWV, MTW, LSP, GAS, GMT, and HEVD performed tissue preparation, sectioning, antibody stainings, confocal microscopy, and analyzed imaging data. SLG and NSG performed behavioral assays and analyzed behavioral data. ECO conceived the project, designed experiments, analyzed data, wrote the manuscript, and acquired funding.

## MATERIALS AND METHODS

### Animals

Adult wild-type zebrafish (*Danio rerio*) of both sexes were kept and bred in a filtered aquarium system (Aquaneering, San Diego, CA) located at Hope College’s zebrafish facility. The aquarium room was kept on a 12-hr light: 12-hr dark cycle at 28° C. Fish were fed 3 times a day *ad libitum*, with commercial flake food (Aquaneering, San Diego, CA) and freshly hatched brine shrimp (Brine Shrimp Direct, Ogden UT) twice and once a day, respectively. All experiments were carried out in accordance with the National Institutes of Health Guide for the Care and Use of Laboratory Animals (NIH Publication No. 80-23). Animal use and research protocols were approved by the Institutional Animal Care and Use Committees from Hope College (HCACUC). All efforts were made to minimize the number of animals used.

### Intracerebroventricular (i.c.v.) 6-OHDA injections

To ablate dopaminergic neurons in the olfactory bulb, we performed intracerebroventricular (i.c.v.) injections of 6-hydroxydopamine (6-OHDA; Sigma-Aldrich, St. Louis, MO) in the dorsal telencephalic ventricle, at the junction of the olfactory bulb with the telencephalon. For this, fish were anesthetized by submersion in a cooled 0.03% MS222 (tricaine) solution (Sigma-Aldrich, St. Louis, MO) until the fish were unresponsive to a tail pinch. We used a beveled syringe with a 26-gauge needle (Hamilton, Reno, NV) inserted diagonally through the skull into the telencephalic ventricle to inject 0.5 μl of a 10 mM of a 6-OHDA solution (Evans blue 1% w/v in PBS evans blue; Sigma-Aldrich, St. Louis, MO) at a rate of 0.01 μL/sec using an Aladdin syringe pump (World Precision Instruments, Worcester, MA). We determined injection success by observing the diffusion of the injected Evans blue solution within the ventricular system. Fish were left to recover for either 1-, 3-, or 7 days post-injection (dpi). Fish from the pranlukast (PRAN; Sigma-Aldrich, St. Louis, MO) treatment received an injection of 0.5 μl of a solution containing 10 mM 6-OHDA with 10 mg/ml pranlukast in Evans blue 1% w/v in PBS. SHAM fish were injected with 1% Evans blue in PBS. PRAN and SHAM fish were sacrificed 1 dpi. Recovered fish were euthanized by over-anesthetization with tricaine, decapitated, and fixed with 4% paraformaldehyde (PFA; Sigma-Aldrich, St. Louis, MO) in PBS for 24 hours at 4°C. The next day, we carefully dissected brains and olfactory organs and prepared the tissue for immunohistochemistry.

### Tissue processing

#### Sectioned tissue for immunohistochemistry

After fixation, we incubated tissue in solutions of increasing ethanol concentrations to dehydrate it, followed by an incubation in xylene (Sigma-Aldrich, St. Louis, MO). Dehydrated tissue was embedded in paraffin (Paraplast plus, McCormick Scientific, Berkeley, CA) and rapidly cooled to solidification. The next day, we obtained semi-serial sagittal 10-μm sections that were then adhered to charged slides (ThermoFisher, Waltham, MA).

#### Cell proliferation assays

To determine cell proliferation, we performed a pulse-and-chase assay with the thymidine analog, 5-bromo-2′-deoxyuridine (BrdU, Sigma Aldrich). We administered 10 μL of a 50 mg/ml BrdU solution in PBS intraperitoneally (i.p.) immediately after the i.c.v. 6-OHDA injection. Fish then were allowed to recover as described above.

### Immunohistochemistry

#### Sectioned tissue preparations

We prepared tissue for immunohistochemistry (IHC) by rehydrating it through descending ethanol solutions followed by antigen retrieval with a 10 mM sodium citrate pH 6.0 solution (Sigma-Aldrich, Canada) at 100°C for 10 min. Next, we blocked slides with a buffer containing 3% normal goat serum (NGS; Vector Laboratories, Burlingame, CA) and 0.4% Triton X-100 (Sigma Aldrich, St. Louis, MO) for at least 1 hour at RT or overnight at 4°C. Next, slides were incubated with primary antibodies (Table 1) overnight at RT, washed, and then incubated with fluorescently-labeled secondary antibodies for up to 2 hours (Table 1). We added 1 μg/ml of 4’,6-diamidino-2-phenylindole (DAPI; BD Pharmingen, Franklin Lakes, NJ) to the secondary antibody solution as a nuclear counterstain. Next, sections were washed and then coverslipped with PVA-DABCO (Sigma Aldrich, St. Louis, MO). Tissue sections were then examined with a confocal laser-scanning microscope Nikon A1 using NIS-Elements software (Nikon, Japan).

**Table 1.** Antibodies used in the study.

| Antibody | Species | Dilution | Source |
| --- | --- | --- | --- |
| HuC/D | Mouse | 1:100 | ThermoFisher |
| GFAP | Rabbit | 1:500 | DAKO |
| TH | Rabbit | 1:500 | Millipore |
| BrdU | Rat | 1:100 | Abcam |
| SV2 | Mouse | 1:1000 | DSHB |
| LCP1 | Rabbit | 1:100 | Genetex |
| Secondaries anti-mouse and rabbit IgG | Goat | 1:200 | Invitrogen |

#### Densitometry

We utilized Adobe Photoshop (Adobe, Mountain View, CA) to assess the optical density (O.D.) of stained tissue with HuC/D and Lcp1. We quantified and averaged 3 to 5 tissue sections per animal. We used images taken at 20x magnification and converted to 8-bit gray scale to obtain whole-section mean luminosity values, which were then converted to OD by the following formula: OD =-log (intensity of background/intensity of area of interest).

#### Cell quantification

Dopaminergic TH (tyrosine hydroxylase), Lcp1+, BrdU+, and TUNEL+ somata were manually quantified. We used 4 to 6 sections of images taken at 20x magnification per animal, and quantified and averaged cells per section.

#### Olfactory-mediated behavioral assays

We analyzed swimming behavioral responses to two odorants, alanine and cadaverine. We used different behavioral chambers to better study different swimming parameters. In all cases, fish were fasted for 48 hours in individual tanks before the experiment and were placed individually in the behavioral chambers. A trial was defined as 30 minutes of general acclimation to the chamber, followed by a 30-minute silent acclimation period. After both acclimation periods elapsed, a 30-second baseline recording was taken prior to odorant delivery. Then, an odorant solution and PBS were administered simultaneously on opposite sides of the chamber. Recording continued for 30 seconds following addition of the compounds and zebrafish behavior was analyzed. After each trial, fish were transferred to another chamber with fresh water and left to acclimate for another trial. Fish underwent 3 or 4 behavioral trials per day. To minimize the effect of the experimenters’ presence, the chambers were surrounded by white panels with perforations for the odorant tubes. The odorant tube was kept the same to control for possible leftover solution in the next trial, but the side of the odorant delivery was randomized for each trial.

For studying behavioral responses to cadaverine, fish were placed in rectangular clear tanks (24.8 cm x 9.7 cm x 15.9 cm) outfitted with two surgical tubes on opposite sides. The chambers were filled with 1.5 liters of fresh fish water before each trial. We used syringes attached to the tubes to administer 1 mL of a 100 μM cadaverine solution (Sigma-Aldrich) in PBS and vehicle (PBS) in the opposite tube. We used a digital camera positioned in front of the tank and behind a white panel to minimize the effect of its presence on the fish.

For testing responses to alanine, we used a white cylindrical behavioral apparatus with 30 cm diameter outfitted with two surgical tubes found on opposite sides about 8 cm above the base. This chamber was filled with 2.5 liters of fresh fish water before each experiment. We used syringes attached to the tubes to administer 3 mL of a 100 μM alanine solution (Sigma-Aldrich, St. Louis, Mo) in PBS and vehicle (PBS) in the opposite tube. We used an overhead digital camera positioned 1 m above to capture swimming behavior (Calvo-Ochoa et al, 2025).

We generated anosmic fish by anesthetizing fish in tricaine as described above. Using a fine paintbrush, we applied cyanoacrylate glue on top of both nasal cavities to temporarily occlude the nares.

#### Video Analysis

All the swimming behaviors we assessed correspond to well-characterized, stereotypical behaviors described extensively in the literature and by our group (Kalueff et al., 2013; Calvo-Ochoa et al. 2025).

For cadaverine analyses, we coded time (in seconds) spent darting and freezing. Darting was characterized as a sudden increased speed of movement and rapid, sharp directional changes. Freezing was characterized by an absence of movement, often accompanied by sinking to the bottom of the tank. We calculated the percent change of response in pre- and post-trials using the formula: percent change= (pre-trial value – post-trial value) / post-trial value x 100 or the percent time spent in erratic swimming or freezing.

For alanine, we used ToxTrac [64], an animal tracking software, to analyze swimming behavior recordings after exposure. Analyses were divided into the 30 seconds of pre-odorant delivery and the 30 seconds of post-odorant delivery. Each recording was uploaded to iMovie in order to increase the contrast to allow for better detection on ToxTrac. Swimming distances (mm) were obtained from ToxTrac.

## Statistical analyses

Comparisons between groups were carried out using Analysis of Variance (ANOVA) with Tukey’s post hoc tests. p-values less than 0.05 were considered significant. Behavioral responses to alanine were analyzed using a one-Sample Wilcoxon test with a baseline of 100% to compare the percentage change of response among groups. All statistical analyses were performed using GraphPad Prism Software (GraphPad, San Diego, CA).

## Notes

### Competing Interest Statement

The authors have declared no competing interest.

